# Principal Metabolic Flux Mode Analysis

**DOI:** 10.1101/163055

**Authors:** Sahely Bhadra, Peter Blomberg, Sandra Castillo, Juho Rousu

## Abstract

**Motivation:** In the analysis of metabolism using omics data, two distinct and complementary approaches are frequently used: Principal component analysis (PCA) and Stoichiometric flux analysis. PCA is able to capture the main modes of variability in a set of experiments and does not make many prior assumptions about the data, but does not inherently take into account the flux mode structure of metabolism. Stoichiometric flux analysis methods, such as Flux Balance Analysis (FBA) and Elementary Mode Analysis, on the other hand, produce results that are readily interpretable in terms of metabolic flux modes, however, they are not best suited for exploratory analysis on a large set of samples.

**Results:** We propose a new methodology for the analysis of metabolism, called Principal Metabolic Flux Mode Analysis (PMFA), which marries the PCA and Stoichiometric flux analysis approaches in an elegant regularized optimization framework. In short, the method incorporates a variance maximization objective form PCA coupled with a Stoichiometric regularizer, which penalizes projections that are far from any flux modes of the network. For interpretability, we also introduce a sparse variant of PMFA that favours flux modes that contain a small number of reactions. Our experiments demonstrate the versatility and capabilities of our methodology.

**Availability:** Matlab software for PMFA and SPMFA is available in https://github.com/ aalto-ics-kepaco/PMFA.

**Supplementary information:** Detailed results are in Supplementary files. Supplementary data are available at https://github.com/aalto-ics-kepaco/PMFA/blob/master/Results.zip.

## Introduction

Principal component analysis (PCA) is one of the most frequently applied statistical methods in Systems Biology (Ma and Dai, 2011; Yao *et al.*, 2012; Barrett *et al.*, 2009). PCA is used to reduce the dimensionality of the data while retaining most of the variation in the data-set (Shlens, 2014). This reduction is done by identifying directions, i.e. linear combination of variables, called principal components, along which the variation in the data is maximal. By using a few such components, each sample can be represented by relatively few variables compared to thousands of features. It also helps us to distinguish between biologically relevant variables and noise.

In the context of transcriptomics and fluxomics, PCA has been widely applied (Yao *et al.*, 2012; Barrett *et al.*, 2009), where a principal component (PC) identifies linear combinations of genes or enzymatic reactions whose activity changes explain a maximal fraction of variance within the set of samples under analysis. The main goals of PCA in fluxomic data are (i) to identify which parts of the metabolism retain the main variability in flux data and (ii) to relate them to the samples, i.e, behaviour of the organism for particular experimental condition.

However, in the context of fluxomics, PCA has some limitations (Folch-Fortuny *et al.*, 2016). PCA considers reactions independently without considering any other structure or relationship among reactions, including stoichiometric relations implied by metabolic pathways. PCA simply extracts a set of reactions that are important to describe sample variance. Moreover, the principal components output by PCA are known to be generally dense, thus including most of the variables, which precludes their interpretation of pathways of any kind. It would be more useful for modelling and biological interpretation if the sample variance captured by the model could be expressed in terms of metabolic pathways or flux modes.

In this paper we propose a novel method to find metabolic flux modes that explains the variance in gene expression or fluxomic data collected from heterogeneous environmental conditions without requiring a fixed set of predefined pathways to be given. The proposed method is called as principal metabolic flux mode analysis (PMFA). Here each principal component, called *principal metabolic flux mode* (PMF), is found by selecting a set of reactions which represents a metabolic flux mode which is approximately in steady state and explains most of the data variability. In addition, we propose a sparse variant, called Sparse Principal Metabolic Flux Mode analysis (SPMFA), to further help the interpretation of the principal components.

Our method differs from existing methods in the literature such as Flux Balance Analysis (FBA) (Orth *et al.*, 2010) as well as more recent proposals as our method aims to explain the sample variability, while existing methods aim to extract flux modes that maximize an objective such as growth as in FBA, or a dominant flux modes active in a set of samples (von Stosch *et al.*, 2016; Folch-Fortuny *et al.*, 2016). Related to our approach, Folch-Fortuny *et al.* (2015) has previously proposed multivariate curve resolution-alternating least squares to improve the biological interpretation of the principal components. Their method incorporates a few constraints such as non-negativity and selectivity when constructing the output. In addition, their method requires a fixed set of metabolic pathways to be defined as a initial step. Very recently, the Principal Elementary Mode Analysis (PEMA) was proposed (von Stosch *et al.*, 2016; Folch-Fortuny *et al.*, 2016) where each component or principal elementary mode are selected from the complete set of elementary modes (EMs) (Pey and Planes, 2014) of the metabolic network such that the selected EMs are responsible for expression levels in a global data. This method needs to derive all possible elementary flux modes explicitly which prevents it to be applicable to genome-scale networks. Moreover, Folch-Fortuny *et al.* (2016, 2015) considered that all fluxes are in steady state, which restricts the applicability of the method in experiments containing transients or perturbations (Baxter *et al.*, 2007).

We verify our proposed PMFA and SPMFA methods by both simulation data and real world fluxomic and transcriptomic data-sets. We demonstrate our model’s flexibility in the analysis of both steady-state and transient data by a change of one regularization parameter. We further show that SPMFA is effective in capturing flux models in experiments involving whole-genome networks.

## 2. Methods

### 2.1 Basic methods

Here we shortly review the existing basic methods for the analysis of fluxomic data.

#### Principal component analysis

We assume 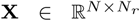 be the data matrix of flux of *N* samples and *N*_*r*_ reactions, with each entry corresponding to a estimated reaction rate for a particular reaction in a particular experiment. We assume that all variables have been centered to have zero empirical mean. The empirical covariance matrix is then given by 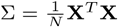 Denoting ∑_1_ = ∑, the 1^*st*^ principal component (PC) w_1_ can be found by solving

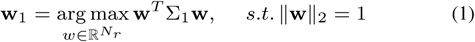

Above 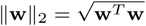 is the *l*_2_ norm of the vector w. The second PC can be found by applying Eq.(1) on updated the covariance matrix using deflation as 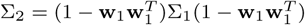 (Mackey, 2009).

The weights, also called the loadings, of the principal component 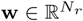 can be interpreted as the importance of reactions in explaining the variance in fluxomic data. The principal components are generally dense, containing most of the reactions of the metabolic network. Sparse PCA (Zou *et al.*, 2006) aims to increase the interpretabilty of PCA by finding principal components that have a small number of non-zero weights through solving the following optimization problem

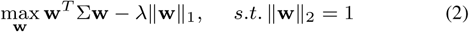

where *λ* is a user defined hyper-parameter which control the degree of sparsity on PC. However, the principal components extracted by neither method represent metabolic flux modes, and will not in general adhere to thermodynamic constraints on reaction directions.

#### Stoichiometric modelling

The metabolic balance of the metabolic system is described using the exchange Stoichiometric matrix 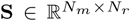 (Raman and Chandra, 2009) which contains transport reactions for inflow of nutrients and output flow of products, but does not contain any external metabolites (as they cannot be balanced). Rows of this matrix represent the *N*_*m*_ internal metabolites, columns present the *N*_*r*_ metabolic reactions including transport reactions and each element S_*m,r*_ shows participation of the *m*^*th*^ metabolite in the *r*^*th*^ reaction: S_*m,r*_ = 1 ( or −1) indicates that reaction *r* produces (or consumes) the metabolite *m*. The value S_*m,r*_ = 0 indicates metabolite *m* is not involved in the reaction *r*. For a flux vector **w**, **Sw** gives the change of metabolic concentration for all metabolites. The metabolic steady-state is assured by imposing a constraint Sw = 0.

#### Elementary modes

The concept of an elementary mode (EM) (Pey and Planes, 2014) is key for the analysis of metabolic networks. An EM is defined as a minimal set of cellular reactions able to operate at the steady-state, with each reaction weighted by the relative flux that they need to carry for the mode to function. An EM also satisfies the reaction directionality constraints arising from thermodynamics.

#### Flux balance analysis (FBA)

FBA (Orth *et al.*, 2010) finds steady state flux modes maximizing objective function. Typically, FBA is done with an objective of maximizing biomass production by solving following optimization problem

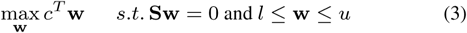

Here *c* indicates the row from Stoichiometric matrix corresponding to biomass production.

### 2.2 Principal Metabolic Flux Mode Analysis (PMFA)

Here we describe our approach, Principal Metabolic Flux Mode Analysis (PMFA), that combines the PCA and Stoichiometric modelling views of metabolism.

To obtain meaningful solutions of steady state flux distributions as PC loading one can impose two additional constraints in PCA formulation:

(1) the weights associated with irreversible reactions should always be positive, i.e., *w*_*ir*_ *≥* 0, where *ir* is an index of an irreversible reaction.

(2) System is in a steady state, where the internal metabolite concentrations do not change, i.e. the metabolite producing and consuming fluxes cancel each other out: Sw = 0.

Considering (1) and (2) the modified optimization problem for doing PCA with structural constraint is as following

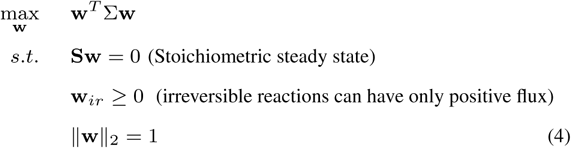

The constraint ║w║_2_ = 1 restricts the spurious scaling up of the weights in the solution. Here, Sw = 0 is a hard constraint and may impose too much restriction when the data does not actually arise from steady-state conditions, e.g. given transients or perturbations of the fluxes during the experiment. Numerically one needs to solve a set of linear equation of size *N*_*M*_ *× N*_*R*_ which makes the problem also computationally hard to solve Eq.(4). Hence instead of considering this hard constraint on the PC loadings we introduce a soft constraint which penalizes the deviation from the steady state. Our aim is to find a flux which optimizes a combination of (1) maximal explained sample variance W^*T*^ ∑W and (2) minimal deviation from a steady-state condition, expressed in the *l*_2_ norm: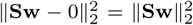. This entails solving the following optimization problem:

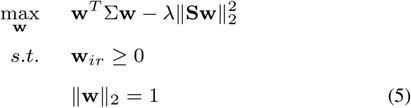

Here *λ* imposes the degree of hardness of the steady-state constraint. For *λ* = 0 the Eq.(5) produces loadings similar to PCA with the exception of the reaction directionality constraint. The model will be henceforth denoted as PMFA(^*l*2^). If desirable, we can make our model to disregard reaction directionality simply by dropping the inequality constraints *w*_*ir*_ *>* 0. We call this version of the method as rev-PMFA or reversible-PMFA.

The *l*_2_ norm on **Sw** in Eq.(5) has the tendency to penalize large steady state deviations in individual reactions, at the cost of favoring small deviations in many reactions. This is probably the desired behaviour in case the data comes from conditions where there is no subsystems that is considerably farther from steady state than other parts of the system. In order to capture the opposite scenario, where a small subset of reactions have large deviation from steady state, one can use *l*_1_ norm regularizer on Sw. The *l*_1_ norm regularizer ║Sw║_1_ in Eq.(5) puts the emphasis of pushing most of the steady-state deviations to zero, whilst allowing a few outliers, metabolites that markedly deviate from steady state. Using *l*_1_ regularizer and a trade-off parameter *λ* we get to solve the following optimization problem:

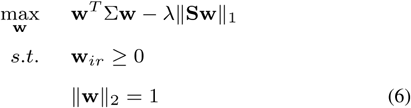

Here *λ* imposes the degree of hardness of the steady-state constraint. Similarly to Eq.(5) for *λ* = 0 the Eq.(6) also produces loadings similar to PCA with selective non-negative constraint. The model will be hence forth denoted as PMFA (*l*_1_).

### 2.3 Sparse principal metabolic flux model analysis

The above formulation of PCA with Stoichiometric constraint still suffers from the fact that each principal component is a linear combination of all possible reaction activities, thus it is often difficult to interpret the results. This problem can be avoided by a variant of PMFA, the sparse principal metabolic flux mode analysis (SPMFA) using an *l*_1_ regularizer (Tibshirani, 1996) on **w** to produce modified principal components with sparse loadings.

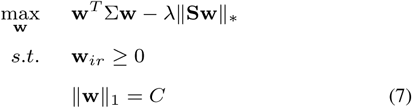

where *║·║*_*\**_ can be any of the *l*_2_ and *l*_1_ norm and *C* is a used defined hyper-parameter which controls the degree of sparsity in principal metabolic flux (PMF) loadings. Similarly to PMFA, Sparse PMFA can also be made consider all reaction reversible by dropping the inequality constraints *w*_*ir*_ *≥* 0. We call this variant rev-SPMFA.

Again, similar to PCA, PMFA aims at explaining the main variability in data using a few PCs. If the original variables have strongly different means and/or variances, the PCs may focus on explaining only the variables with the highest values and/or variances, disregarding the small variance associated with the rest of variables. Hence before applying PMFA we need to centralize the expression and fluxomic data.

### 2.4 Algorithms

The objective function of Eq.(5) can be interpreted as difference of two differentiable convex function. This type of optimization problem is known as Difference of Convex functions (DC) program. We used the convex-concave procedure (CPP), a local heuristic that utilizes the tools of convex optimization to find local optima of difference of convex functions (DC) programming problems (Lipp and Boyd, 2016). Using CCP method we solved Eq.(5) by solving following convex approximation (quadratic program) in each iteration *t*:

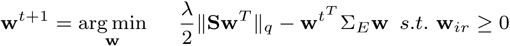

followed by projecting w^*t*+1^ on *║w║*_*p*_ = *C*. The norms *p, q ∈* {1, 2} are chosen according to the desired model.

To obtain a *multi-factor* PMFA model, i.e. a model containing several PMFs jointly representing the data, we follow a approach similar to some PCA algorithms, namely the deflation of the covariance matrix. However, due to additional stoichiometric constraint here we deal with a sequence of non-orthogonal vectors, [**w**_1_*,…,* **w**_*d*_] hence we must take care to distinguish between the variance explained by a vector and the additional variance explained, given all previous vectors. We have used orthogonal projection for deflating the data matrix (Mackey, 2009). This also maintain positive definiteness of covariance. For every iteration *d*+1 we first transfer already found principal flux modes 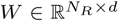 to a set of orthogonal vectors, {*q*_1_*,…, q*_*d*_}.

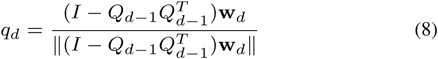

where, *q*_1_ = w_1_, and *q*_1_*,…, q*_*d*_ form the columns of *Q*_*d*_. *q*_1_*,…, q*_*d*_ form an orthonormal basis for the space spanned by w_1_*,…,* w_*d*_. Then the Schur complement deflation of covariance matrix is done by

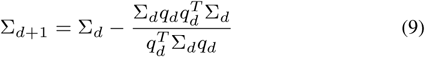

## 3. Results

We report a comparative study on following methods.

- PCA: Principal component analysis as given by Eq.(1). PCA_*dir*_ denotes the PCA augmented with reaction directionality constraints.
- SPCA: Sparse PCA corresponding to Eq.(2). SPCA_*dir*_ is the SPCA augmented with reaction directionality constraints.
- FBA: Flux balance analysis with an objective of maximizing biomass production given by (3).
- PMFA: Principal Flux Mode Analysis as described in Section 2.2. 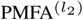 denotes *l*_2_ regularization on the stoichiometric constraint Eq.(5) while 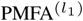 denotes *l*_1_ regularization on stoichiometric constraint Eq.(6).
- SPMFA: Sparse Principal Flux Mode Analysis as given by Eq.(7). Again, 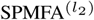 denotes *l*_2_ regularization on stoichiometric constraint, while 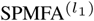 denotes *l*_1_ regularization on stoichiometric constraint.
- Principal Elementary Mode Analysis (PEMA) (von Stosch *et al.*, 2016; Folch-Fortuny *et al.*, 2016): It uses the set of EMs as the candidates for the PCs. It models the flux matrix **X** is as follows:

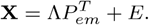

Above, *P*_*em*_ is the *N*_*r*_ *× N*_*f*_ principal elementary mode matrix, formed by a subset of *N*_*f*_ EMs from the entire EM matrix; Λ is the is the *N × N*_*f*_ non negative weighting matrix; and *E* is the *N × N*_*r*_ residual matrix. *P*_*em*_ is found by iteratively selecting important EMs. We only used PEMA on small metabolic networks since as calculation of all EMs for genome-scale metabolic networks is impractically time consuming (Pey and Planes, 2014).

### Data centralization

PCA, SPCA, PMFA, and SPMFA aim at explaining the main variability in data using a few PCs. If the original variables have strongly different means and/or variances, the PCs may focus on explaining only the variables with the highest values and/or variances, disregarding the small variance associated with the rest of variables. Hence before applying all of them, we need to centralize the expression and fluxomic data.

### Selection of optimal level of regularization

We selected the optimum levels of the regularization parameter *λ* for PMFA and SPMFA and level of sparsity for SPMFA by cross-validation maximizing the *fraction of sample variance captured* on test samples

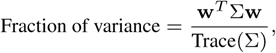

which is a classic measure used with PCA and related approaches. Above, w is the PC computed from the training data, and ∑ is the co-variance matrix of the test sample. Leave-One-Out (LOO) cross-validation was used on smaller datasets and 5-fold cross-validation was used on the large whole genome dataset.

### 3.1 Datasets

#### Pichia pastoris simulation case study

This case is based on the metabolic network of Pichia pastoris, which originates from Tortajada *et al.* (2010).

It describes the central carbon metabolism of P. pastoris during growth on glucose, glycerol and methanol, comprising the Embden-Meyerhoff-Parnas pathway, citric acid cycle, penthose phosphate and fermentation pathways. It contains 45 compounds (36 of which are internal metabolites, which can be balanced for growth) and 44 reactions, yielding a total number of 98 EMs (Tortajada *et al.*, 2010). Flux data were generated simulating the growth of Pichia pastoris for twelve different cultivation conditions by choosing appropriate sets of active EMs. The active EMs were assumed to contribute randomly to the flux pattern. Hence we can compare PMF identified by PMFA to the *ground truth* “active EMs” that were used for data generation.This case study also enables the study of the impact of noise on the EMs identification and performance. For this study we add random Gaussian noise to fluxomic data, where noise variances are 2%, 5%, 10% and 20% of original values. From the flux data and the deviation reported in supplementary material of Quek *et al.* (2009) we observed that most the reported fluxes have deviation associated with it and the deviations are in rage of 2-5% of their reported value along with few reactions with deviations even more than 12% of their value.

#### Saccharomyces cerevisiae experimental case study

A metabolic network for Saccharomyces cerevisiae proposed by Hayakawa *et al.* (2015) and fluxome data used in (von Stosch *et al.*, 2016; Hayakawa *et al.*, 2015; Frick and Wittmann, 2005) was used in this study. The network describes the central cytosolic and mitochondrial metabolism of S. cerevisiaecomprising glycolysis, the pentose phosphate pathway, anaplerotic carboxylation, fermentative pathways, the TCA cycle, malic enzyme and anabolic reactions from intermediary metabolites into anabolism (von Stosch *et al.*, 2016). The network contains 42 compounds (30 of which are internal metabolites, which can be balanced for growth) and 47 reactions. The objective in this case study is to evaluate the performance of PMFA Eq.(5) on fluxome data and compare it with PEMA and PCA. For PEMA we have used 1182 EMs provided by von Stosch *et al.* (2016).

#### Saccharomyces cerevisiae gene expression data

The objective of experiment described in this section is to evaluate the performance of the proposed PMFA Eq.(5) and SPMFA Eq.(7) on transcriptomic data and compare it with PCA and SPCA.

For studying applicability of the proposed PMFA (Eq.(5)) in transcriptomic data where system may not be in steady state, we considered transcriptomic data both in steady state as well as time series. The steady state transcriptomic data has been generated by Rintala *et al.* (2009) where Saccharomyces cerevisiae grown in glucose-limited chemostat culture with 0%, 0.5%, 1.0%, 2.8%, or 20.9% O2 in the inlet gas (D= 0.10 /h, pH5, 30C) (Wiebe *et al.*, 2008). The normalized transcription data-set is available in the Gene Expression Omnibus (GEO) database (Barrett *et al.*, 2011) with the accession number GSE12442. It contains four samples for 0,0.5,2.8 and 20.9% oxygen and six samples for 1% oxygen and samples were either from four (0,1,20.9% oxygen) or two independent cultivations (0.5, 2.8% oxygen). This data-set is combined with time-series transcriptomic data generated by Rintala *et al.* (2011) where time series analysis starting from two (1% and 20.9%) levels of oxygen provision. Seven time points at 0, 0.2, 3, 8, 16, 24, 72*/*79 hours from both time series and two biological replicates from each time point were analysed The microarray data can be accessed through GEO accession number GSE22832 (Barrett *et al.*, 2011). We have used Yeast community model v. 7.5 ( YCM 7.5), which contains 3494 reactions among 2220 compound and catalysed by 909 genes. We have converted gene expression data to a expression level per reaction by the method described in the method section.

Given a gene expression data we have converted it to reaction vs. sample data-set using the following gene rules. Let us denote *X*^*G*^ as gene expression matrix with size *N × N*_*G*_ where *N*_*G*_ is number of genes and the *g*^*th*^ column of 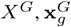 is the expression vector corresponding to gene *g*. Then,

- if gene association with reaction *r* is denoted as *g*_1_ **or** *g*_2_ then expression value for reaction *r*, i.e. *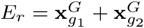.*
- otherwise if gene association with reaction *r* is denoted as *g*_1_ **and** *g*_2_ then expression value for reaction *r*, i.e. *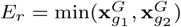.*

### 3.2 Prediction of active EMs using PFMA

In our first experiment we evaluated the predictive performance the proposed PMFA and PEMA in correctly retrieving underlying active elementary flux modes. We used the Pichia pastoris simulation case study data, where the elementary flux modes that are part of the ground truth are known. For the evaluation, area under ROC curve (AUC) and area under precision recall curve (AUPR). The precision/recall metrics, widely used in information retrieval, is to assess how well the flux modes computed by PEMA and PMFA correlate with the ground truth active EMs.

For each PMF, we computed its correlation with respect to all 98 elementary flux modes of the Pichia pastoris metabolic network. We then sort the EMs in descending order of correlation and consider first *i* = 1 *…,* 98 EMs as the predicted EMs by the model. Precision and recall is then computed for each *i*, by considering ground truth active EMs within the first *i* EMs as true positives and other EMs with the top *i* as false positives. A precision/recall curve can be then plotted by taking the precision/recall values for all *i*s, in the order of the descending correlation in the sorted list. The AUPR is denoted as area under the precision recall curve and AUC is denoted as area under receiver operating characteristic curves (Hanley and McNeil, 1983).

In a multi-factor PMFA model, to compute a precision recall value for a *k*-factor model we considered the maximum correlation of an EM with any of the *k* factors as a final correlation of an EMs with multi-factor PMFA model. Then, we sorted all EMs according to descending order of their maximum correlations. With PEMA model we used an analogous approach: for a *k*-factor PEMA, for each *i* we included the top *i* correlated EMs (according to the maximum correlation of EMs with any of *k* factors) as the models prediction and used those for computing the precision/recall values for each *i* = 1*,…,* 98.

Figure 1 shows total AUPR and total AUC achieved by the different models for different amount of additional noise. It shows that PMFA is robust with respect to noise in the fluxomic data, with both AUPR and AUC metrics only slowly decreasing as a function of increasing noise, until noise level of 10%. In this regime, adding more factors to PMFA models also increases performance monotonically both in AUC and AUPR metrics, showing that the additional factors recover EMs that were not captured by the first factor. In the high noise regime (*>* 10%) we observe that the performance of the 3-factor PMFA model drops suggesting that the last factor likely starts to capture noise.

**Fig. 1.**
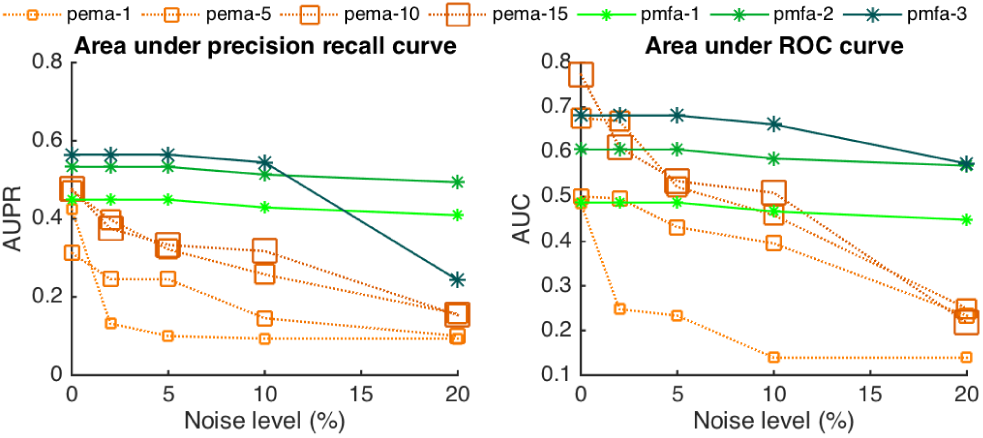
The graph shows the AUPR (left) and AUC(right) values obtained by different models for different noise levels.

In the noise free case, PEMA performs comparatively to PMFA, especially in terms of the AUC metric and when using a high enough number of factors in the model. However, the performance of PEMA deteriorates quickly upon increased noise. The decrease of performance is particularly apparent in the AUPR metric. Moreover, we note that the PEMA models do not exhibit as clear monotonic improvement when adding more and more factors to the model as PMFA, as in Figure 1 the performance curves of the different PEMA models cross each other in several occasions.

### 3.3 Explaining test set variance with PMFA

In this experiment we focused on the ability of PMFA to explain variance on data in a predictive setting, that is, on new data that has not been used for model estimation. We focused on the amount of variance explained in the test set in a Leave-One-Out (LOO) cross-validation setting.

We studied the effect of stoichiometric regularization 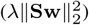 on the fraction of sample variance captured by PMFA and alternative models (PEMA, PCA). Figure 2 shows the fraction of sample variance explained by the first PMFs and PCs as a function of deviation from steady state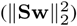 in test data of two fluxomic data-sets (S. cerevisiae and P. pastoris). The deviation from the steady-state is controlled by the regularization parameter *λ ≥* 0: high values of *λ* give low deviation from steady-state and vice-versa.

**Fig. 2.**
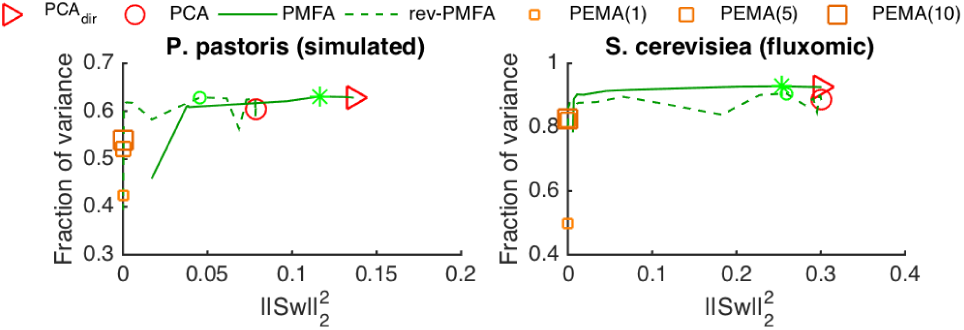
Depicted is for two fluxomic data-sets the fraction of variance on test data in LOO setting as a function of deviation from steady state 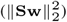 captured by PCA, directional PCA (PCA_*dir*_), 1-, 5-and 10-factor PEMA, as well as PMFA and rev-PMFA using different amount of Stoicihiometric regularization. The markers ‘*\**’ and ‘_*o*_’ indicate the optimal level of regularisation for PMFA and rev-PMFA.

In particular on the fluxomic datasets, relatively heavy regularization can be applied without decrease of variance explained, indicating that the data can be well explained by steady-state flux modes.

By change of the regularisation parameter *λ*, the statistics of PMFA exhibit a continuous transition from fully steady state flux modes 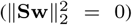i.e. PCA augmented with reaction directionality constraints. The transition for rev-PMFA is not as smooth as PMFA with directionality constraint. It is evident that the directionality constraint increases the stability of PMFA without reducing much explained variance on test data.

Compared to PEMA, The fraction of variance explained the first PMF from rev-PMFA is higher than 1-, 5-and 10-factor PEMA regardless of the amount of stoichiometric regularization or application of the directionality constraints. The amount of variance explained by the first PMF from PMFA is also much higher than 1-factor PEMA even with high Stoichiometric regularization, while the 5-and 10-factor PEMA reach the level of PMFA for both data sets.

Figure 3 shows the explained fraction of variance on test data in a Leave-One-Out (LOO) cross-validation setting, where both test and training data is contaminated with various amount of the noise. The test set variance captured by first component of PMFA only very slightly decreases upon increasing noise. In contrast, the test set variance captured by PEMA drops considerably when the noise level increases. Higher order PEMA models are here somewhat more resistant than the 1-factor PEMA but still not competitive with PMFA. In addition, we note that PCA is not able to explain test set variance as well as PMFA regardless of the noise level. To understand this result, we note that within the training set, by definition we expect PCA to explain the variance the best. However, when analysing new data not seen in the training phase, the stoichiometric information used by PMFA helps to attain a superior predictive performance.

**Fig. 3.**
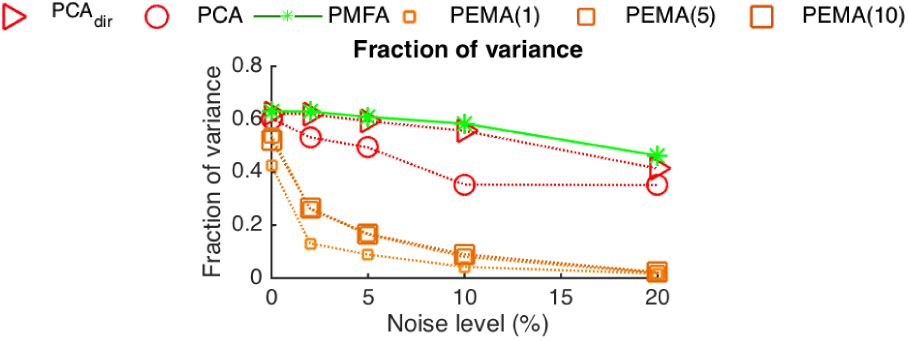
Depicted is for the P. pastoris simulated data-set the fraction of variance on test data in LOO setting as a function of additional noise level captured by PCA, PCA_*dir*_ 1-, 5-and 10-factor PEMA, as well as PMFA (with optimum regularization parameter).

### 3.4 Recovery of sparse flux modes from full genome data by SPMFA

In this experiment, we evaluated the Sparse Principal Metabolic Flux Mode Analysis, SPMFA, in discovery of sparse flux modes. We focus on the full genome data, i.e., all steady-state and transient samples of *S. cerevisiae* containing a total of 3494 reactions for, making dense principal components and flux modes difficult to interpret. To quantify the fraction of explained variance normalized by the complexity of the extracted flux mode, we measure the *normalized fraction of variance*, calculated as

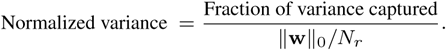

Above, ║w║_0_ denotes the *l*_0_ norm, i.e. the cardinality of non-zero elements of w Figure 4 shows variance (left) and normalized variance (right) as the function of deviation from steady state 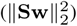

**Fig. 4.**
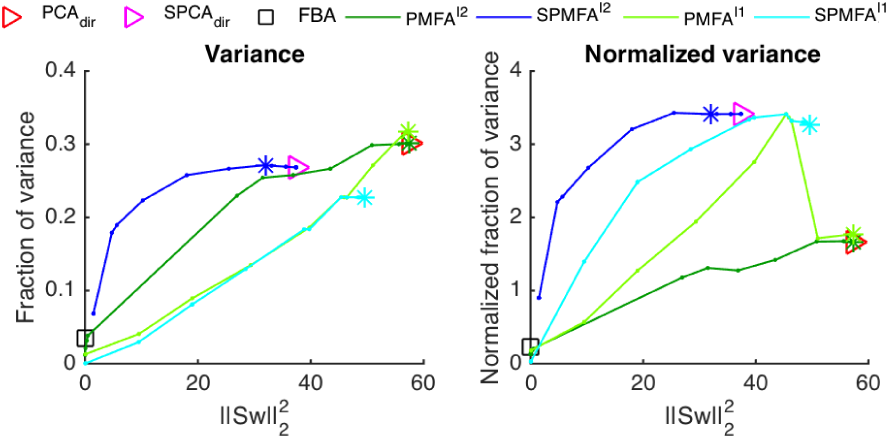
Figure shows variance (left) and normalized variance (right) on test data in 5 fold cross validation setting as a function of steady state deviation 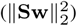 on the whole genome gene expression data (containing both steady-state and transient samples) for PMFA SPMFA and FBA. The markers ‘*\**’ indicate the optimal level of regularisation.

At the maximum, PMFA captures slightly more explained variation than SPMFA at (Figure 4, left). Correspondingly, SPMFA is vastly more effective in capturing normalized variance, achieving more than double the rate of PMFA at any level of deviation from steady state ((Figure 4, right). SPMFA statistics can be seen to smoothly approach the (directional) sparse PCA statistics when the deviation from steady-state is let to increase.

The variant 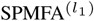 which is regularized by the *l*_1_ stoichiometric regularizer (║Sw║_1_), also exhibit a smooth transition, but captures less variance at the maximum, albeit the fraction of normalized variance at the maximm,albeit the fraction of normalized varience captured is similar to SPMFA. 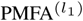 exhibits a phase change, following PMFA at high steady state distances (small *λ*) but switching to SPMFA regime as regularization is increased. This reflects the fact that with small *λ* the model is not yet sparse but sparsity quickly emerges once *λ* is increased.

It is notable that on this large heterogeneous dataset, all methods fail to capture meaningful amounts of normalized sample variance in the vicinity the steady state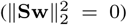 This is also true for FBA, which we have included as a comparison (maximum biomass production as the FBA objective). The FBA solution is sparse but the fraction of variance captured is very small, causing as the normalized variance captured by FBA to be small compared to SPMFA solution when the stoichiometric regularization is relaxed. This illustrates the importance of being able to relax the steady-state assumption when analyzing real-world experiments.

### 3.5 Analysis of SPMFA on S. cerevisiae oxygen series gene expression data-set

In this section we analyze the principal flux modes found by SPMFA(_*l*_2) (Eq.(7)) separately on gene expression data of (a) transient time-series samples and (b) steady state samples (c.f. Section 3.1) on a whole-genome metabolic network of S. cerevisiae. In section 3.3, we have observed that both PMFA and SPMFA explain similar amount of sample variance but SPMFA is vastly more effective in terms of normalized fraction of variance. We have also found that, each of the first few PMFs selects more than thousand reactions, while each SPMF has only around 300 active reactions. For better interpretation we select SPMFA for this study.

Henceforth, we will denote the first three components of SPMFA for time-series data-set as T1, T2 and T3 respectively. Similarly, the first three components of SPMFA for steady-state data-set will be denoted as SS1, SS2 and SS3. We have found that the principal components T3 (time-series SPMF3) and SS3 (steady-state SPMF3) are effectively identical and contain only reactions pertaining to the cell envelope, which is a well-known response to variable oxygen levels. We also observe that many reactions are common in different SPMFs of time-series and steady-state data-set. Figure 5 shows a Venn diagram for set of the reactions selected by different flux modes. The sparse principal metabolic flux mode T2 (time-series SPMF2) and SS1 (steady-state SPMF1) have 167 reactions in common and again the sparse principal metabolic flux mode T1 (time-series SPMF1) and SS2 (steady-state SPMF2) have 210 reactions in common.

**Fig. 5.**
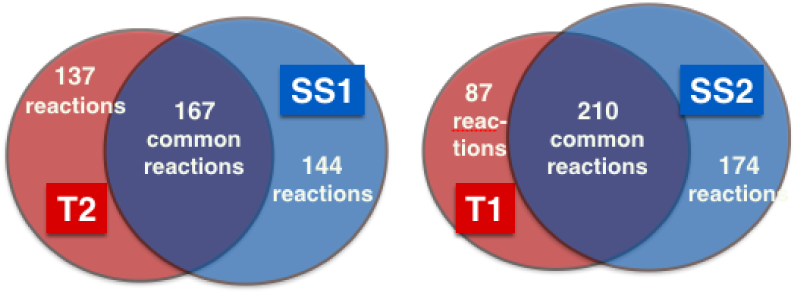
Figure shows a Venn-diagram of reactions selected by SS1(28.45% variance captured),SS2(8.67%),T1(28.34%) and T2(1.45%). It shows that 176 reactions are common in SS1 and T2 while 210 reactions are common in SS2 and T1.

Furthermore, the four principal components have a few reactions in common. Therefore, we define seven metabolic *modes* from four SPMFs. Table 1 presents a description of these seven metabolic modes. The reactions in Mode 1 included cytosolic NADPH-production by the oxidative pentose phosphate pathway, NADPH-consuming dehydrogenases, and NADH-transport from cytosol to mitochondria via the malate-shuttle. Mode 2 represents down-regulation of many sterol transporters, and up-regulatation aldehyde dehydrogenases and ribonucleotide reductases for the biosynthesis of DNA nucleotides. Mode 3 contains many biosynthetic pathways such as chorismate, heme, glutamate, nucleotide, and glycogen biosynthesis. It also contains sugar transporters, lower glycolysis, oxidative TCA cycle, and several tRNA loading enzymes. Therefore, Mode 3 is strongly associated with growth. Mode 4 contains biosynthetic pathways for spermidine, acyl-carrier protein (ACP), and phosphoribosyl diphosphate (PRPP). PRPP is a precursor for histidine and nucleotides. Mode 4 represents up-regulation of thiamine, glycerol, and nucleotide transporters. It also picks up upregulation peroxisomal enzymes, tRNA loading enzymes, and enzymes in lower glycolysis. Mode 4 can thus be considered growth with the peroxisome active. Mode 5 contains biotin synthesis, triose phosphate isomerase, and transporters for serine, aspartate, glutamate, phosphate, glycine, and ammonia. Mode 5 is unique in that upregulation/downregulation directions which are opposite to that in Mode 3 and Mode 4. Mode 7 contains biosynthesis pathways for shikimate and isoprenoids, NAD translocation to mitochondria, and a shuttle for NADPH from cytosol to mitochondria via aldehyde dehydrogenases.

**Table 1.** Description of the seven metabolic flux modes we recovered from four SPMFs of both the data-sets, i.e., T1 and T2 for time-series data-set and SS1 and SS2 for steady-state data-set. It also contains the flux mode of T3 or SS3.

| Subset | Content |
| --- | --- |
| Mode 1: Reactions only in T1 | PPP, malate-shuttle |
| Mode 2: Reactions only in T2 | sterol transporters, deoxyribonucleotides |
| Mode 3: Reactions in T1 and SS2 | TCA, glycogen, heme, nucleotides, and translation = growth |
| Mode 4: Reactions in T2 and SS1 | Peroxisome, spermidine, thiamine, ACP and translation = growth |
| Mode 5: Reactions in T1, T2, SS1 and SS2 | Biotin synthesis and nitrogen reallocation transporters |
| Mode 6: Reactions only in SS1 | - |
| Mode 7: Reactions only in SS2 | Shikimate, isoprenoids, NAD translocation, and NADPH shuttle |
| Reactions in T3 and SS3 | cell envelope |

## 4. Discussion

In this paper we have proposed a novel method for the analysis of metabolic networks, called the Principal Metabolic Flux Analysis, PMFA, through the combination of stoichiometric flux analysis and principal component analysis, finds flux modes that explain most of the variation in fluxes in a set of samples. Unlike most stoichiometric modeling methods, PMFA is not tied to the steady-state assumption, but can automatically adapt— by the change of a single regularization parameter—to deviations from the stoichiometric steady-state, whether they are due to measurement errors, biological variation, or other causes. Our experiments showed that the method is more robust to the steady-state violations than competing approaches, and can compactly capture the variation in the data by a few factors. For the analysis of whole-genome metabolic networks, we further proposed Sparse Principal Flux Mode Analysis, SPMFA that allows us to discover flux modes with a small fraction of reactions activated, thus could be interpreted as pathways. Our experiments showed that our methods are more efficient in capturing the variance in sets of experiments than methods based on elementary flux mode analysis or flux balance analysis. The efficient Concave Convex Procedure optimization allows the method to scale up to whole-genome models unlike methods based on search in the space of elementary flux modes.

Analysis of cultivation data on the whole-genome metabolic network of Saccharomyces cerevisiae showed that SPMFA was able to identify many pathways responsive to changes in oxygenation, including well-known pathways such as cell envelope, PPP, TCA, glycolysis, sterol transport, hexose transport, heme, nucleotides, peroxisome, spermidine, thiamine, biotin, shikimate, and isoprenoid pathways. In addition, the analysis grouped these pathways in new subsets which may lead to novel insight. In particular, the existence of two growth modes; one with lower and one with higher peroxisomal activity, is interesting from an engineering point of view when pathways utilizing peroxisomal proteins are used in novel synthetic pathways. This analysis reveals a potentially novel shuttle for NADPH from cytosol to mitochondria via aldehyde dehydrogenases.

## Acknowledgements

We acknowledge Merja Oja and Eija Vartiainen for their suggestions and comments and the computational resources provided by the Aalto Science-IT project.

## References

Barrett, C. L., Herrgard, M. J., and Palsson, B. (2009). Decomposing complex reaction networks using random sampling, principal component analysis and basis rotation. BMC systems biology, 3(1), 30.

Barrett, T., Troup, D. B., Wilhite, S. E., Ledoux, P., Evangelista, C., Kim, I. F., Tomashevsky, M., Marshall, K. A., Phillippy, K. H., Sherman, P. M., et al. (2011). Ncbi geo: archive for functional genomics data sets–10 years on. Nucleic acids research, 39(suppl 1), D1005–D1010.

Baxter, C., Liu, J., Fernie, A., and Sweetlove, L. (2007). Determination of metabolic fluxes in a non-steady-state system. Phytochemistry, 68(16), 2313–2319.

Folch-Fortuny, A., Tortajada, M., Prats-Montalbán, J. M., Llaneras, F., Picó, J., and Ferrer, A. (2015). Mcr-als on metabolic networks: Obtaining more meaningful pathways. Chemometrics and Intelligent Laboratory Systems, 142, 293–303.

Folch-Fortuny, A., Marques, R., Isidro, I. A., Oliveira, R., and Ferrer, A. (2016). Principal elementary mode analysis. Molecular BioSystems, 12(3), 737–746.

Frick, O. and Wittmann, C. (2005). Characterization of the metabolic shift between oxidative and fermentative growth in saccharomyces cerevisiae by comparative 13 c flux analysis. Microbial cell factories, 4(1), 1.

Hanley, J. A. and McNeil, B. J. (1983). A method of comparing the areas under receiver operating characteristic curves derived from the same cases. Radiology, 148(3), 839–843.

Hayakawa, K., Kajihata, S., Matsuda, F., and Shimizu, H. (2015). 13 c-metabolic flux analysis in s-adenosyl-l-methionine production by saccharomyces cerevisiae. Journal of bioscience and bioengineering, 120(5), 532–538.

Lipp, T. and Boyd, S. (2016). Variations and extension of the convex–concave procedure. Optimization and Engineering, 17(2), 263–287.

Ma, S. and Dai, Y. (2011). Principal component analysis based methods in bioinformatics studies. Briefings in bioinformatics, 12(6), 714–722.

Mackey, L. W. (2009). Deflation methods for sparse pca. In Advances in neural information processing systems, pages 1017–1024.

Orth, J. D., Thiele, I., and Palsson, B. Ø. (2010). What is flux balance analysis? Nature biotechnology, 28(3), 245–248.

Pey, J. and Planes, F. J. (2014). Direct calculation of elementary flux modes satisfying several biological constraints in genome-scale metabolic networks. Bioinformatics, page btu193.

Quek, L.-E., Wittmann, C., Nielsen, L. K., and Krömer, J. O. (2009). Openflux: efficient modelling software for 13 c-based metabolic flux analysis. Microbial cell factories, 8(1), 25.

Raman, K. and Chandra, N. (2009). Flux balance analysis of biological systems: applications and challenges. Briefings in bioinformatics, 10(4), 435–449.

Rintala, E., Toivari, M., Pitkänen, J.-P., Wiebe, M. G., Ruohonen, L., and Penttilä, M. (2009). Low oxygen levels as a trigger for enhancement of respiratory metabolism in saccharomyces cerevisiae. BMC genomics, 10(1), 1.

Rintala, E., Jouhten, P., Toivari, M., Wiebe, M. G., Maaheimo, H., Penttilä, M., and Ruohonen, L. (2011). Transcriptional responses of saccharomyces cerevisiae to shift from respiratory and respirofermentative to fully fermentative metabolism. Omics: a journal of integrative biology, 15(7-8), 461–476.

Shlens, J. (2014). A tutorial on principal component analysis. arXiv preprint arXiv:1404.1100.

Tibshirani, R. (1996). Regression shrinkage and selection via the lasso. Journal of the Royal Statistical Society. Series B (Methodological), pages 267–288.

Tortajada, M., Llaneras, F., and Picó, J. (2010). Validation of a constraint-based model of pichia pastoris metabolism under data scarcity. BMC systems biology, 4(1), 1.

von Stosch, M., de Azevedo, C. R., Luis, M., de Azevedo, S. F., and Oliveira, R. (2016). A principal components method constrained by elementary flux modes: analysis of flux data sets. BMC bioinformatics, 17(1), 1.

Wiebe, M. G., Rintala, E., Tamminen, A., Simolin, H., Salusjärvi, L., Toivari, M., Kokkonen, J. T., Kiuru, J., Ketola, R. A., Jouhten, P., et al. (2008). Central carbon metabolism of saccharomyces cerevisiae in anaerobic, oxygen-limited and fully aerobic steady-state conditions and following a shift to anaerobic conditions. FEMS Yeast Research, 8(1), 140–154.

Yao, F., Coquery, J., and Lê Cao, K.-A. (2012). Independent principal component analysis for biologically meaningful dimension reduction of large biological data sets. BMC bioinformatics, 13(1), 1.

Zou, H., Hastie, T., and Tibshirani, R. (2006). Sparse principal component analysis. Journal of computational and graphical statistics, 15(2), 265–286.

